# Tailored Photoacoustic Apertures with Superimposed Optical Holograms

**DOI:** 10.1101/2023.08.01.551479

**Authors:** Glenn A. Howe, Meng-Xing Tang, Christopher J. Rowlands

## Abstract

A new method of generating potentially arbitrary photoacoustic wavefronts with optical holograms is presented. This method uses nanosecond laser pulses at 1064 nm that are split into four time-delayed components by means of a configurable multipass optical delay apparatus which serves to map the pulses onto phase-delayed regions of a given acoustic wavefront. A single spatial light modulator generates separate holograms for each component which are imaged onto a photoacoustic transducer comprised of a thermoelastic polymer. The spatially- and temporally-modulated holograms facilitate a phased array effect which enables beam steering of the resulting acoustic pulse. For a first experimental demonstration of the method, as verified by simulation, an acoustic beam is steered in four directions by around 5 degrees.

## 1. Introduction

Ultrasound has been an invaluable tool for medical imaging for decades, and the properties of which have been studied extensively [1, 2, 3]. However, even as enlightening as the modality has proven to be, there exist numerous limitations such as the static configuration and number of transducer elements [4, 2, 5], and operator nuances leading to potentially inaccurate or inconsistent measurements [6, 7]. Furthermore, conventional ultrasound transducers are typically arrays of elements numbering on the order of hundreds for linear arrays to thousands for 2D arrays [2, 5]. This is orders of magnitude smaller than the number of elements in typical optical imaging systems, which commonly employ megapixel-scale resolutions.

Since photoacoustics introduces an optical-to-acoustic process regarding the generation of the ultrasound pulse [8, 9, 10], especially in the case of using laser pulses to irradiate a thermoelastic polymer, this modality employs some flexibility not seen in traditional ultrasonics. Photoacoustic ultrasound owes this flexibility to the fact that the transducer ‘elements’ can be arbitrarily arranged in both shape and number due to the optically-based nature of the irradiation patterns. Indeed, a photoacoustic transducer can theoretically approach a continuous ‘array’, much like how an optical aperture can be considered continuous. Careful modulation of both the temporal and spatial extent of this irradiation, as described in this study, can lead to wide ranging versatility of acoustic wave generation, resulting in photoacoustic spatial ultrasound modulation–conceptually a configurable acoustic phased array.

To produce high-fidelity optical patterns comprising the high-resolution photoacoustic aperture, the incident light is first modulated by refractive, diffractive, or other optical methods. While this may be done by physical elements [11, 12], this technique is largely inflexible since the patterns are static. The use of patterned holograms, by means of spatial light modulators (SLMs), has also shown to be an effective method of modifying acoustic wavefront propagation [13, 14, 15, 16], however this relies on large illumination footprints and often necessitates the use of many acoustic cycles, thus eliminating the axial resolution benefits of single-oscillation pulses. Photoacoustic phase modulation can be carried out by temporal modulation of the incident laser pulses. Previous implementations have used free-space [17, 10] or fiber-based delivery [18, 19, 20, 21, 22, 23, 24, 25, 26, 27, 28], however these methods typically produce static linear phased arrays which are fixed in both number and size. Acoustic phase variation can also be achieved by the use of acoustic lenses which, analogous to optical lenses, directly modulate the incident acoustic pulse [29, 30, 31, 32, 33, 34, 35, 36]. However, these are also static and inflexible in modulating the resulting acoustic wavefront.

Phase control in the photoacoustic aperture plane can alternatively be achieved by mapping the time delay of the incident laser pulses onto the phase delay of the resulting acoustic pulse. In this method, shown below, the acoustic phase is constructed by overlapping the optical patterns of different laser pulses with appropriate intensities, yielding a smooth phase gradient as shown in Fig. 1. The spatial and temporal modulations of the incident optical energy therefore manifest as an arbitrarily-patterned, 2D pulsed phased array.

**Figure 1:**
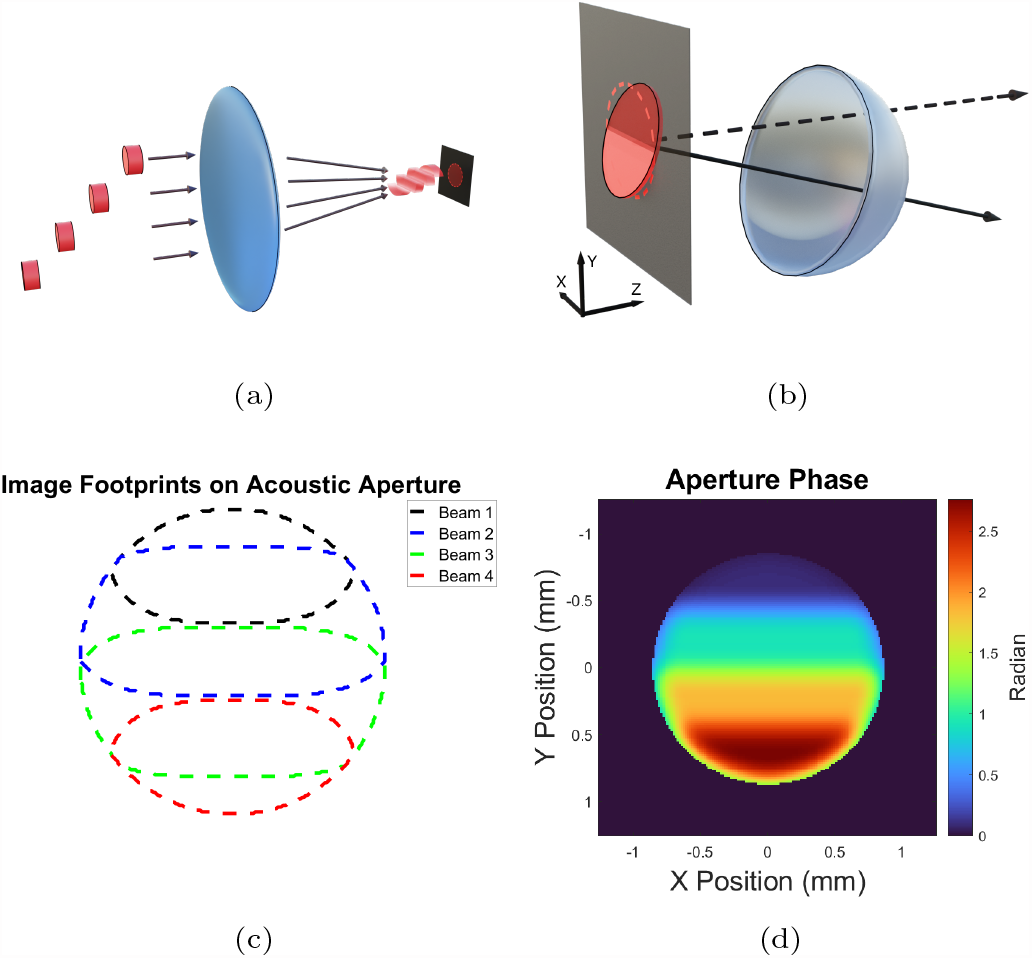
Working principle of the instrument. **(a)** Four time-delayed laser pulses are imaged onto a thermoelastic polymer, creating an acoustic pulse. **(b)** The time delay acts as an acoustic phase delay, resulting in steerable wavefront propagation. **(c)** The optical patterns are modulated and overlapped in such a way that they result in **(d)** a continuous phase ramp across the photoacoustic aperture due to the smooth intensity transitions between individual beam components.

This study seeks to validate the aforementioned acoustic pulse generation concept. As a step towards building an arbitrary acoustic phase in the aperture plane, this approach used four time-delayed pulses to modulate the phase of an acoustic wavefront. A thermoelastic polymer consisting of polydimethylsiloxane (PDMS) and carbon black (CB) was used as a photoacoustic transducer which was irradiated by multiple time-delayed holograms. The holograms, when combined spatially, form an optically-driven acoustic phased array which has the flexibility of creating a wide range of photoacoustic apertures. The design of the optical delay line and description of the experimental setup is provided in Sec. 2, a simulation of expected results is described in Sec. 3, and the experimental results are shown in Sec. 4. A discussion of the results can be found in Sec. 5.

## 2. Methods

### 2.1. Experimental Setup

Described below is the experimental setup with major components described in detail in their own subsection. The working principle of the instrument is illustrated in Fig. 1. Four time-delayed laser pulses, previously patterned by an SLM, were imaged onto a thermoelastic polymer. These SLM patterns, located at the pupil plane of the imaging system, carry optical phase modulations (holograms) that manifest as intensity distributions at the conjugate image plane (i.e. optical images on the photoacoustic transducer). The time-delayed pulses map to acoustic phase delays and therefore steer the resulting acoustic pulse away from normal incidence. The SLM-generated illumination patterns were overlapped in such a manner that the resulting phase ramp in the acoustic aperture is smooth and continuous.

The experimental setup is illustrated in Fig. 2. A nanosecond laser (Litron Nano-L 200-30) emits a single 6 ns, 1064 nm pulse and is immediately expanded from 1 mm diameter to 6.5 mm diameter. This expanded, collimated pulse was then injected into a combined beam splitter/beam delay apparatus which simultaneously splits the single pulse into four components and delays them temporally in succession (Sec. 2.2). These four temporally-delayed pulses were then collinearly aligned in a rectangular pattern before being directed onto the surface of an SLM (Sec. 2.3). The SLM operates on the four pulses independently to produce four distinct diffraction patterns –one for each time-delayed pulse. Still collinearly-aligned, these four diffraction patterns were then imaged onto the surface of a ‘photoacoustic transducer’ which is comprised of a composite thermoelastic polymer (Sec. 2.4).

**Figure 2:**
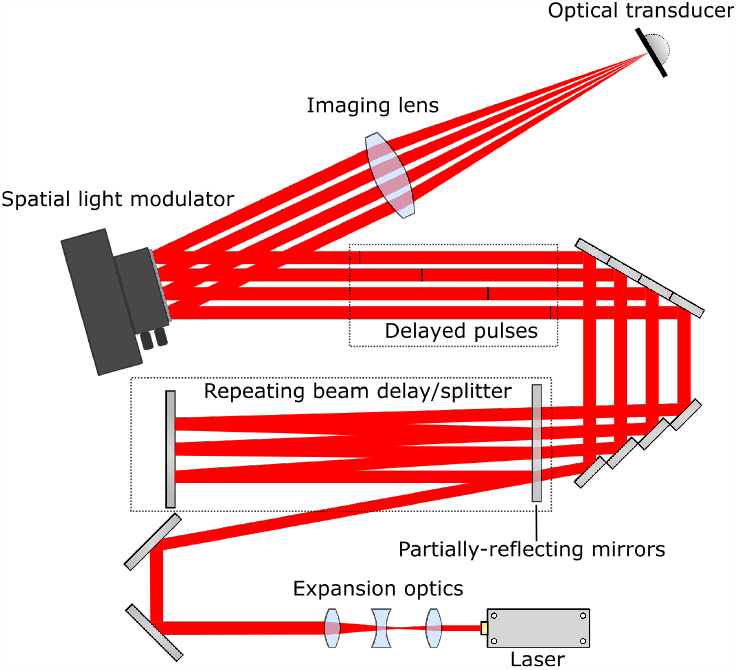
Optical diagram of experimental setup. A single laser pulse is injected into a multipass optical splitter/delay line. The four timedelayed pulses are patterned by a spatial light modulator before being imaged onto a thermoelastic polymer which acts as a photoacoustic transducer. Note that the four beams are collinearly arranged in a rectangular pattern, but shown here in a line for illustrative purposes.

The temporally-delayed patterns (images of the SLM Fourier space) act as sources of laser-generated acoustic pulses, and the time delays of the laser pulses map to the phase of the resulting acoustic pulses. These phase-delayed pulses are used to shape and steer the constructed acoustic wavefront, analogous to a conventional phased array [37]. Measurements of the acoustic wave were captured by a needle hydrophone which raster scans the plane orthogonal to the acoustic wave propagation direction to generate XY slices of the acoustic environment.

### 2.2. Optical Beam Delay

Separating the initial single laser pulse into four was achieved by means of a custom optical delay line. The optical delay apparatus serves to both split the initial laser pulse into four roughly-equal-intensity components as well as to impart a sequential time delay between them as they propagate through the device. Seeing as the temporal delay of the laser pulses map to phase delays of the acoustic pulses, and considering the rapid speed of light, a considerable physical distance is required which necessitated the construction of a custom design. The design uses numerous mirrors to contain the pulses in a multipass loop which allows for a compact footprint while still achieving a large total temporal delay.

The design shares some principles with conventional multipass cells such as the White [38] and Herriot [39] cells in that a beam is temporarily trapped within a reflective cavity. This design, however, uses a number of partially-reflecting mirrors to allow the passage of a controlled amount of light at regularly-spaced intervals. Furthermore, this design allows for a wide range of delays (after reconfiguration and realignment) due to the usage of only flat mirrors. A free-space design was opted for as opposed to a fiber-based design due to considerations relating to both ease of delay tunability as well as laser induced damage when dealing with high power densities. More discussion can be found in the Supplementary Materials, and related diagrams are shown in Fig. 13.

### 2.3. SLM Hologram Generation

Optical pattern generation was carried out via an SLM operating in phase-only mode (Meadowlark HSP1920-500-1200-HSP8). The face of the SLM has a resolution of 1920 x 1152 with a pixel pitch of 9.2 x 9.2 *μ*m. After being temporally-delayed, the four laser pulses were then collinearly aligned in a rectangular pattern which corresponds to four quadrants of the SLM surface, as shown in Fig. 3a. These four quadrants are controlled independently to create four distinct patterns.

**Figure 3:**
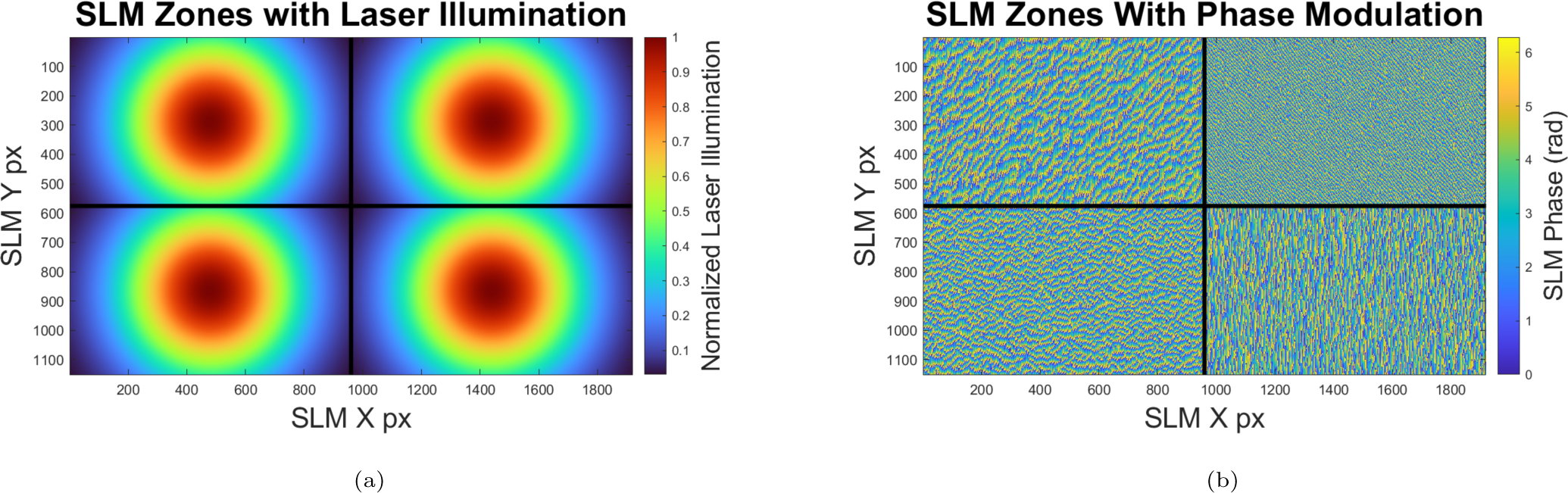
Diagrams of the **(a)** illumination and **(b)** example phase of the SLM surface. The four roughly-equal-intensity beams are collinearly aligned and are operated on by four distinct regions of the SLM surface, allowing for independent control over the four resulting diffraction patterns. The phases shown in (b) are example patterns generated by implementing a Gerchberg-Saxton routine (Section 2.3). The resulting intensity distributions, when imaged through the lens, are shown in Fig. 6.

Pattern generation was carried out by implementing a Gerchberg-Saxton (GS) routine. This is a simple, iterative phase retrieval method which quickly calculates the phase pattern necessary to produce a resulting target image given a known input illumination pattern [40]. For the experiments described here, the GS patterns were calculated and applied to the SLM quadrants prior to data collection, as exemplified in Fig. 3b.

These SLM patterns, along with a blazed grating pattern, allow arbitrary images to be created free of contamination of zero-order diffraction effects. A lens captured the non-zero-order, phase-modulated light and produced the resulting images at the conjugate image plane of the SLM –the surface of the photoacoustic transducer. These images are four temporally-delayed components which were used to construct and shape an acoustic wavefront.

### 2.4. Photoacoustic Transducer

The construction and properties of non-biological optical transducers vary greatly in absorption efficiency, bandwidth, and ease of fabrication [41, 42, 43, 44, 45, 46, 47, 48, 49, 50, 51, 52]. The photoacoustic transducer used in this study was a composite thermoelastic polymer comprised of PDMS and carbon black. Both materials are safe to use, easy to work with, and widely available. The aforementioned studies have investigated the use of similar composites and much work has been done to increase the bandwidth and optical absorption efficiency. Achieving high bandwidth was not a priority in this study. In fact, due to the relationship with the optical delay and acoustic phase, a lower bandwidth is advantageous. Furthermore, due to both the scope of this study and the large overhead in available laser power, maximizing optical absorption was also not a priority.

### 2.5. Needle Hydrophone

To capture the resulting acoustic field generated from the incident laser pulses on the photoacoustic transducer, a needle hydrophone was employed (Precision Acoustics 0.5 mm (NHO500)). This particular device has a working frequency range of 0.2–40 MHz with a typical ±4dB response in the 1–20 MHz range.

Scanning motion of the needle hydrophone was carried out by a microscope stage (Prior Scientific H101A). Its large range of motion (114 x 75 mm) and small step size (0.01 *μ*m) allows both flexibility and repeatability. The stage was oriented vertically enabling lateral, 2D slices of the acoustic field. The hydrophone was affixed to the stage via a custom plate adapter and mounting hardware. Z-axis control was handled by Thorlabs LT1 2-inch manual translation stage.

### 2.6. Data Acquisition

Signals captured from the needle hydrophone were recorded and digitized by a two-channel oscilloscope (Redpitaya STEMlab 125-14). Running at a sampling rate of 125 MHz with a bit depth of 14, the first channel captured the Q-switch trigger signal from the laser while the second channel captured the acoustic signal from the needle hydrophone. A home-built script written in Python handled the collection, parsing, and organization of each measurement while also commanding the vertically-oriented microscope stage to scan a 2D slice of acoustic environment.

## 3. Simulation of Expected Results

### 3.1. Simulation Methods

Traditional phased array wavefront propagation including beam steering and arbitrary wavefront generation can be modeled via simple methods stemming from Huygen’s Principle of wavefront propagation. The Rayleigh-Sommerfeld diffraction (RSD) equation (Eqn. 1) describes the field *u*(*x, y*; *z*) at some downstream z location given the field *u*_0_(*s, t*) at the initial propagation point (*z*_0_):

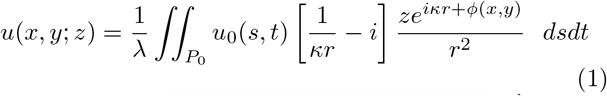

where 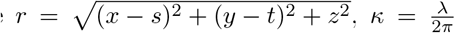, *P*_0_ is the 2D region of the photoacoustic aperture, and *ϕ*(*x, y*) is the local phase offset. The arguments (s,t) and (x,y) describe lateral dimensions in the initial and resulting field, respectively.

While slow, RSD permits the interrogation of all planes of the propagation environment with a simple ‘permutation’ of the Cartesian variables. Furthermore, one may apply an additional envelope to the exponential term, thus permitting an interrogation of the acoustic pulse length. The additional envelope has consequences for beam steering and side lobe formation. While similar results, along with improvements in computation speed, can be achieved using the Angular Spectrum Method (ASM) and related techniques [53], the familiarity and flexibility of RSD propagation outweigh the speed benefits of ASM for this study. The simulations, and analysis thereof, were performed in MATLAB R2021b.

### 3.2. Simulation Parameters

#### 3.2.1. Time and Phase Delay

As described in Sec. 2.2, the sequential time delay between laser pulse components is 36 ns, corresponding to a total delay of 108 ns. Although this time delay is static, it naturally corresponds to different total phase delays for different acoustic pulse wavelengths. Unless stated otherwise, all results in this study map the 108 ns time delay to the given acoustic wavelength in question.

#### 3.2.2. Photoacoustic Aperture

The size of aperture has profound effect on the nature of wavefront formation and beam steering. Not only is diffractive beam spreading a function of aperture size, but also is the overall phase gradient (since the total time delay is constant), and therefore the degree to which a beam can be steered. Prior experimental investigations have guided this study to adopt a photoacoustic aperture diameter of 1.7 mm which has been shown to provide an acceptable optimization between beam spread and steering capabilities. This size regime also affords sufficient optical power density to facilitate an appreciable acoustic pulse amplitude. In other words, the incident SLM-generated laser patterns combine to form an image on the photoacoustic transducer with a total diameter of 1.7 mm.

#### 3.2.3. Simulation Dimensions

Prior experimental investigation had also shown the center frequency of the given thermoelastic polymer to be around 1–10 MHz (dependent upon construction methods and materials used), thus corresponding to a dominant wavelength of roughly 150–1500 *μ*m when assuming a roughly 1500 m/s propagation speed in water. The simulation volume was then taken to be a cube with length of 20 mm which permits investigation of a wide range of transducer apertures while still being well within the Fraunhofer (far-field) regime [54]:

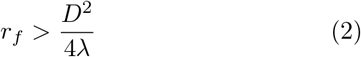

where D is the extent of the radiation source. This allows apertures to be maximally a few mm which is more than adequately large for the beam steering technique laid out in this study.

#### 3.2.4. Simulation Resolution

While the RSD method provides an exact solution at infinitesimal resolution elements [53], an arbitrarily large resolution often yields diminishing returns given the increase in computational expense. To optimize the efficiency and accuracy of the simulation, a given ‘beam forming feature’, in this case beam steering deviation, is measured vs. the simulation grid resolution elements. A simple 108 ns ‘phase’ ramp was applied to an aperture 1.7 mm in diameter, and the resulting beam was propagated to a z distance of 20 mm. Deviations were measured via 2D Gaussian fits to the resulting XY acoustic amplitude profile. Figure 4 shows the results for three acoustic wavelengths spanning 1–10 MHz, highlighting the clear asymptotic trend in each case. Applying a simple power function fit to the data reveals a practical limit to the increase of resolution elements at around 1300 (corresponding to roughly *λ*/10 at 10 MHz), signifying a suitable value that will be used for the remainder of this study.

**Figure 4:**
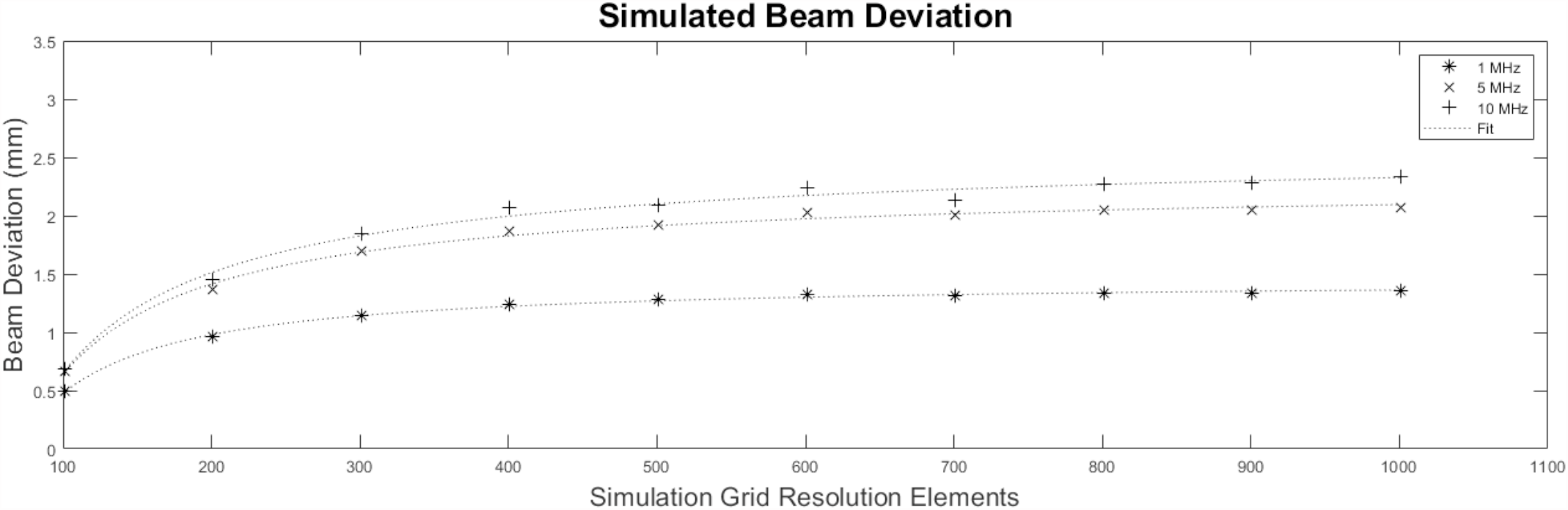
Results of simulating beam steering at constant aperture phase ramp with different simulation resolutions and pulse frequencies. The beam deviations asymptote to 1.4, 2.2 and 2.5 mm for the 1, 5, and 10 MHz frequencies, respectively.

### 3.3. Hologram Generation

A diagram of the SLM surface with the locations of the four collinearly-aligned laser beams is shown in Fig. 3. The SLM contains pre-generated, Fourier-plane patterns which both shape the optical wavefront and also steer the light away from the zero-order diffraction angle. These GS patterns are calculated from the known incident intensity distribution and the intended image plane patterns. An optical lens facilitates the imaging of the four independent patterns simultaneously on a single image plane. Figure 5 shows the intended individual target patterns along with the overlapped pattern as seen by the photoacoustic transducer. Figure 6 shows a recorded image of the target hologram images. The images show a pattern approximately 7 mm in diameter for illustrative purposes, whereas the illumination patterns used in this investigation were 1.7 mm in diameter carrying a total fluence of 15.1 mJ/cm^2^. Speckles are an inherent aspect of discrete hologram imaging, and can be exacerbated by small optical apertures (e.g. laser beam diameters) along with finite Fourier frequency components (e.g. divided SLM surface regions). The effect of the speckle pattern on the resulting acoustic wavefront is an attractive area for investigation, however it remains outside the scope and framework of the current study.

**Figure 5:**
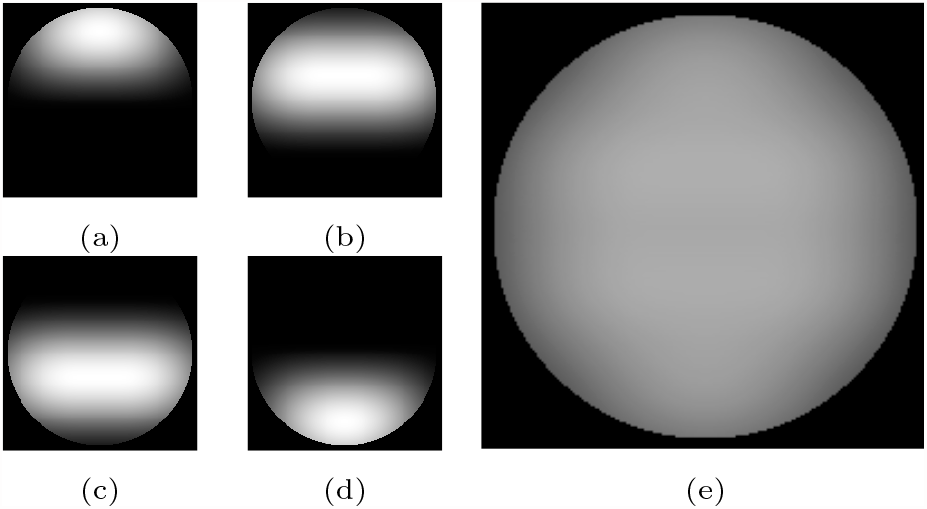
Simulated images of the four SLM hologram patterns as seen by the photoacoustic transducer. When the separate beam patterns **(a-d)** overlap, the result is a complete circular disk **(e)**, thus maintaining an even illumination and smooth phase ramp across the aperture.

**Figure 6:**
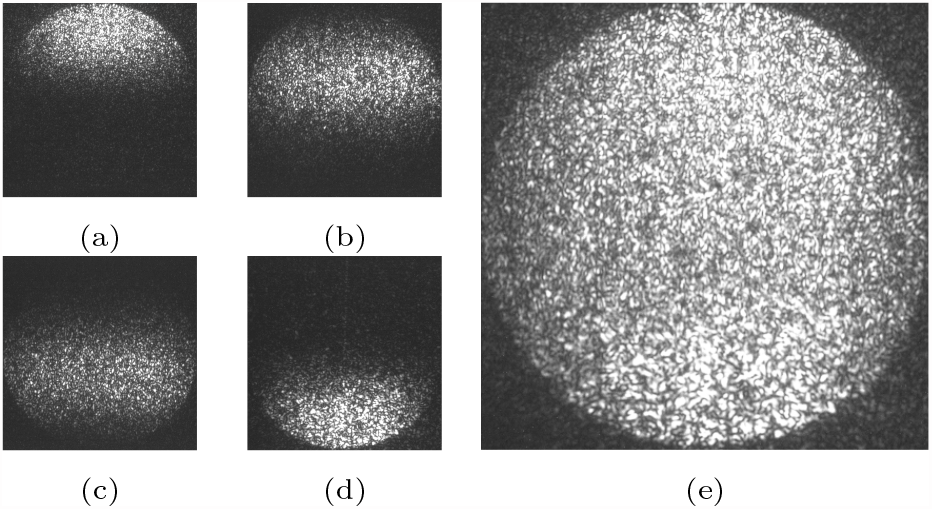
Camera images of the four SLM hologram patterns as seen by the photoacoustic transducer. The individual patterns **(a-d)** were produced by the four separate regions of the SLM and imaged on the photoacoustic transducer. **(e)** shows the images overlapped simultaneously for illustrative purposes when in fact each pulse is sequentially time delayed by around 36 ns.

### 3.4. Acoustic Field Propagation

The photoacoustic aperture intensity and phase distribution was propagated via RSD to a distance of 20 mm using the parameters laid out in the proceeding sections. The resulting XY cross section was downsampled to a resolution of 21 x 21 points, matching the sampling of the needle hydrophone. This acoustic field sampling resolution was found to be an acceptable optimization of data collection speed and wavefront recovery fidelity at the tolerances necessary for this study to confirm conceptual realization of the beam steering method.

Figure 7 shows the results from the RSD wavefront propagation. A flat wavefront at 10 MHz is first shown in Fig. 7a while Figs. 7b to 7d show steered beams with frequencies of 10, 5, and 1 MHz, respectively. The sub panels show, from left to right, the aperture phase, a full-resolution YZ lateral view of beam directionality, a polar plot of beam directionality, and the simulated, downsampled XY transverse hydrophone measurement. Aperture phases were calculated assuming a 108 ns total phase delay for the respective frequencies.

**Figure 7:**
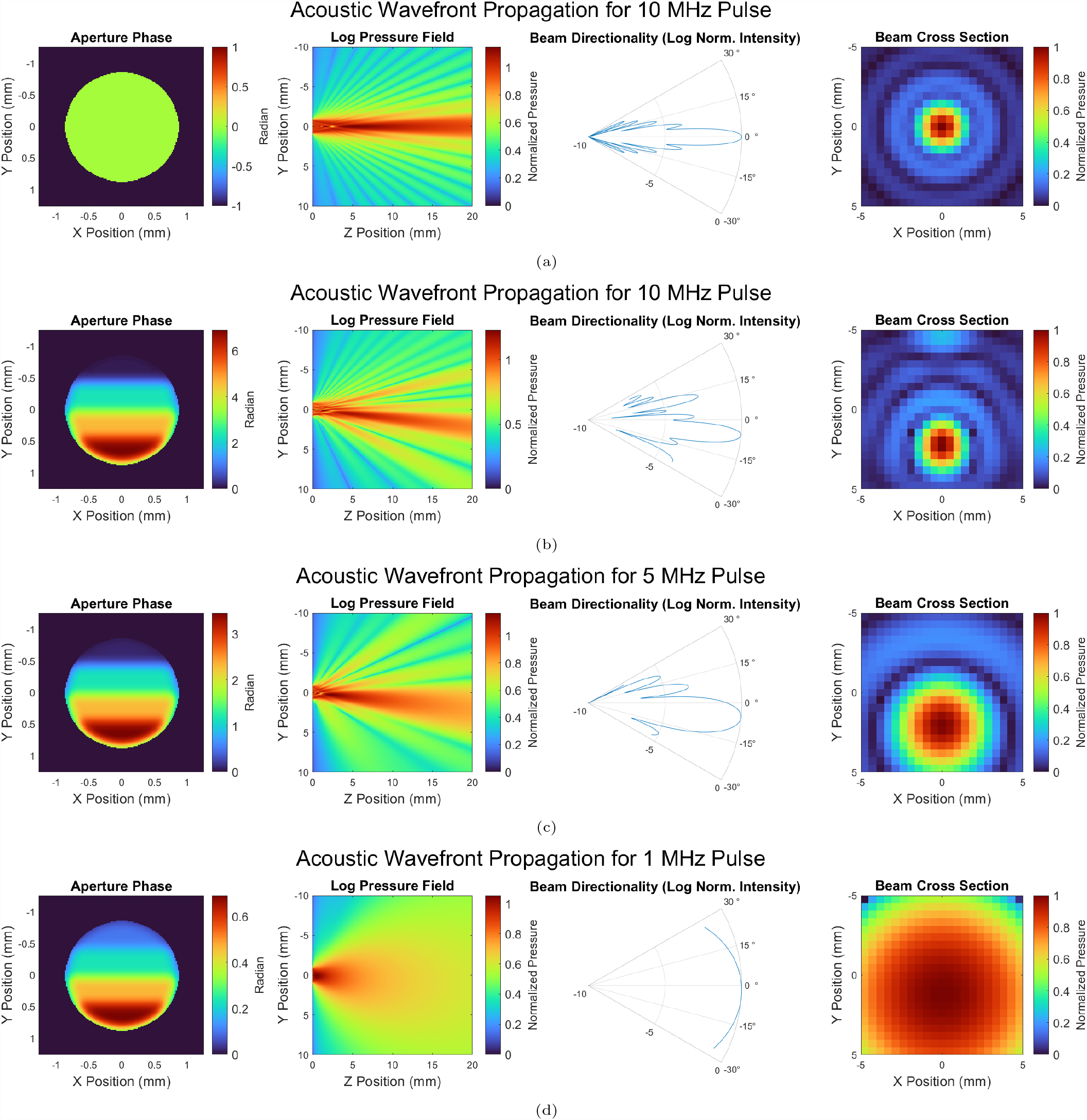
Simulation results of the RSD propagation method for acoustic frequencies ranging from 1–10 MHz. **(a)** shows no beam steering while **(b)-(d)** show smooth phase ramps. In each of (a-d), from left to right, the first panel shows the phase ramp across the aperture, the second shows a YZ cross section (where Z is the propagation direction) of the resulting beam formation, the third shows a corresponding polar plot, and the last shows an XY pressure map at 20 mm. While the total delay was 108 ns for each simulation, this corresponds to different total acoustic phase ramps for different acoustic frequencies.

## 4. Experimental Results

### 4.1. Acoustic Pulses and Bandwidths

Synchronized times series of the four individual acoustic pulse components are shown in Fig. 8, corresponding to the illumination patterns shown in Fig. 6. Pulse intensity normalization was controlled by adjusting the independent diffraction efficiency of each beam within the separate SLM zones shown in Fig. 3. Still, inhomogeneities in the thermoelastic polymer can manifest as modulation of the pulse intensity, as is illustrated by the slightly larger amplitude in the first pulse component of the shown traces. Further discussion can be found in Sec. 5.1

**Figure 8:**
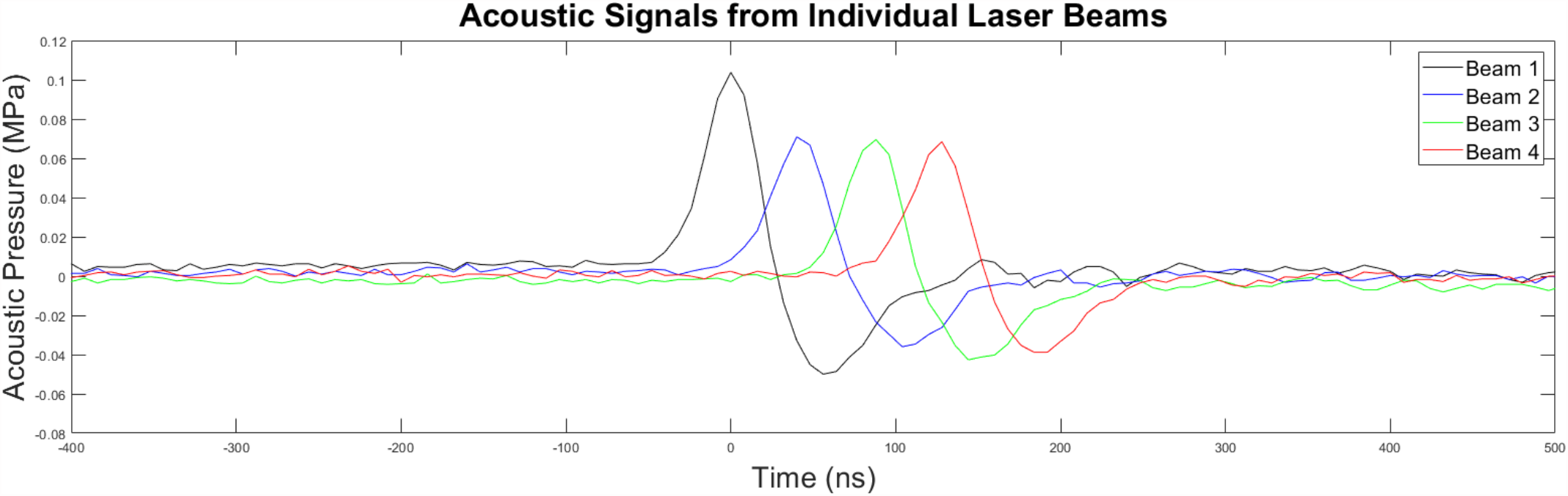
Time series of typical acoustic pulses generated from the individual laser illumination pattern components, synchronized and overlapped. The sequential time delay is around 36 ns. The illumination patterns, as described in Sec. 2.3, spatially overlap in conjunction with the time delay to produce an even phase ramp. Intensity inhomogeneity is likely due to thermoelastic polymer features discussed in Sec. 5.

Previous experiments showed a typical frequency response to a PDMS+CB composite irradiated by a 6-ns 1064 laser to be between 1–10 MHz. For this investigation, the upper limit was targeted to provide adequate bandwidth for both single pulses and multi-component combined pulses which naturally stretch out the emergent pulse period and thus lowers the resulting frequency.

Figure 9a shows a time series and accompanying fast Fourier transform (FFT) of a sample acoustic pulse from a single laser pulse component. As expected, the central frequency lies around 10 MHz. Figure 9b shows similar results for a multi-component combined pulse. It is seen that the central bandwidth shifts towards lower frequencies accompanied by more structure at higher frequencies. This structure is not the governing characteristic of the bandwidth, therefore the dominant frequency is taken to be the maximally-represented frequency of around 4 MHz. The expected results for this particular photoacoustic transducer are assumed to carry a similar dominant frequency.

**Figure 9:**
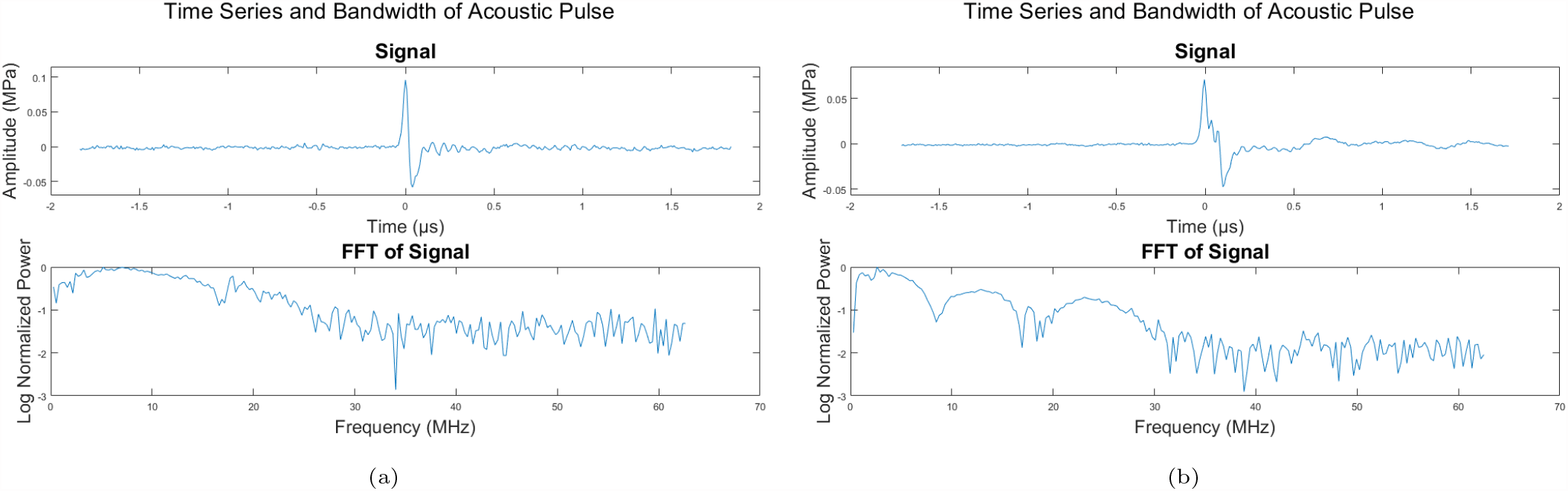
Time series and FFT of examples of a typical **(a)** single and a **(b)** composite pulse. Having been constructed from multiple time-delayed pulses, the composite pulse naturally has a lower peak frequency (along with higher-order structure).

### 4.2. Measured Acoustic Field

Using this 4 MHz as the propagation frequency, along with an aperture diameter of 1.7 mm and total propagation distance of 20 mm, the results of the RSD propagation simulation are shown in Fig. 10. The total expected beam deviation is 2.1 mm corresponding to a beam steering angle of around 5 degrees.

**Figure 10:**
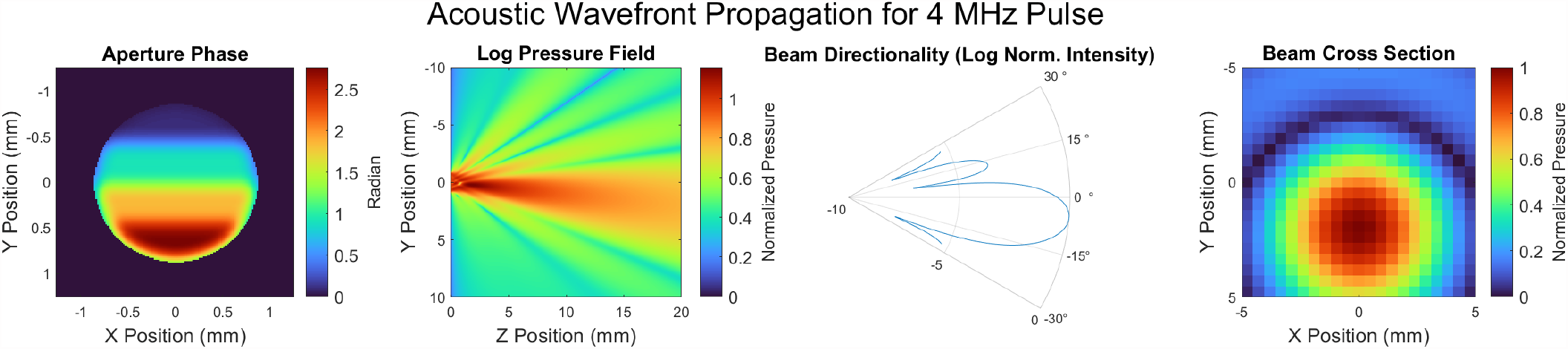
Example of the expected results using the particular photoacoustic transducer described in this study. Given the data shown in Fig. 9, the expected acoustic frequency is 4 MHz and therefore the expected beam deflection after 20 mm is 2.1 mm.

The results of the measured acoustic field are shown in Fig. 11. In the left-hand panels, which show the measured data, an obvious feature not captured by the simulation are the ‘residuals’ residing at the center of each acoustic field. While this feature is discussed in Sec. 5, it is here assumed to be distinct from the steered beams’ amplitude. As such, a 2D Gaussian fit was applied to an exponential threshold of the measured acoustic field to isolate the steered beam from the residuals. These Gaussian fits are shown in the right-hand panels. The black dashed circles indicate the non-steered beam locations while the red dashed circles indicate the expected 2.1 mm deviations as given by the results of the RSD simulation. A summary of the beam steering results is presented in Tab. 1. This table shows the simulated and measured results of the (x, y) deviations for the four cardinal directions, with boldface numbers signifying the intended direction. Three of the four measurements fall within 90% of the expected deviation magnitudes, with minor angular deviations from the expected cardinal directions. The leftward steered beam saw roughly 50% expected deviation and strong zero-order residual.

**Figure 11:**
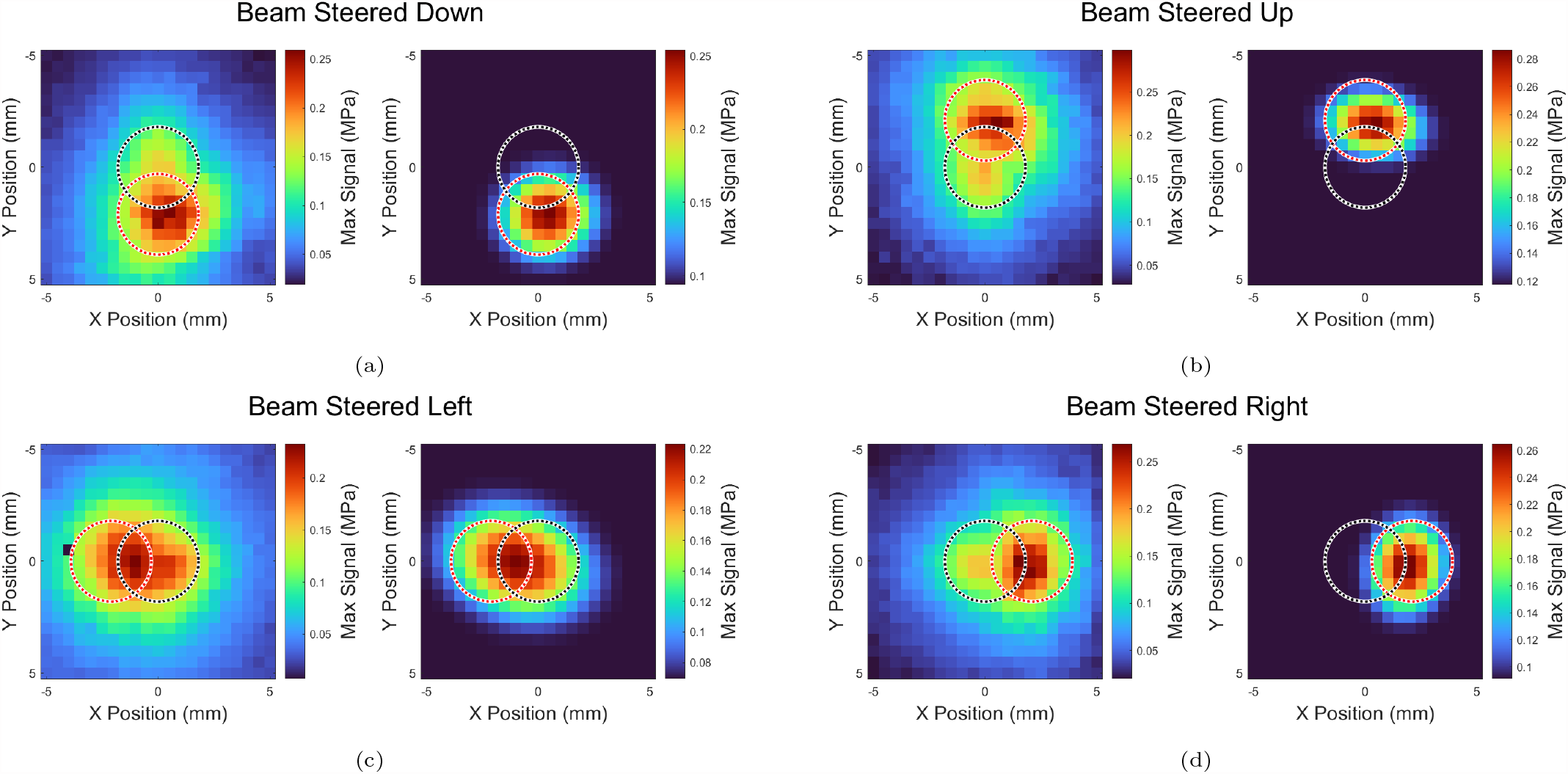
Measured results from the photoacoustic phased array. In each subfigure, the left panel shows the recorded data while the right panel shows Gaussian fits applied to an exponential threshold of the measurements. The non-deviated locations are marked with a black dashed circle while the expected deviations are marked with a red dashed circle. (a), (b), and (d) show *∼*90% agreement with expected results while (c) shows *∼*50% agreement, likely due to thermoelastic polymer inhomogeneity.

As further verification of the beam steering technique, time series at two differing Y-axis locations are shown in Fig. 12a. The earlier wavefront pixel shows a composite nature which is indicative of pulses arriving from the center of the photoacoustic aperture, while the latter wave-front pixel has a pulse profile corresponding to a single pulse component –namely due to the photoacoustic aperture component that is most delayed in phase. The red and blue time series correspond to the red and blue circles in Fig. 12b which shows the acoustic field and corresponding phase map, respectively.

**Figure 12:**
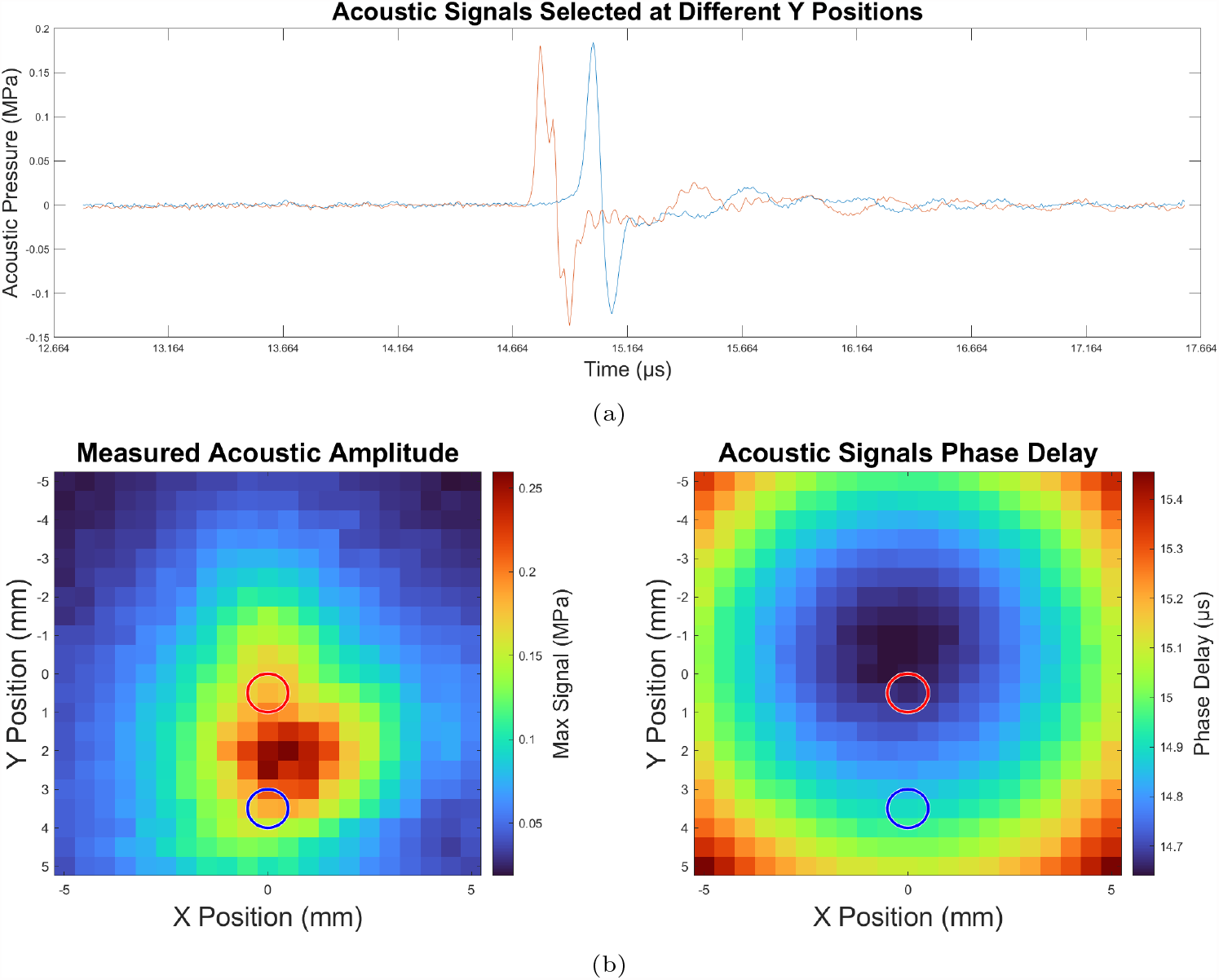
**(a)** Time series of measurements at different Y positions, and **(b)** intensity and phase maps of the measured acoustic field. The red and blue traces in (a) were taken at the locations shown as red and blue circles in (b).

## 5. Discussion and Conclusions

### 5.1. Remarks on Experimental Setup

The residuals, indicated in Sec. 4, have a few likely sources. First, due to the interferometric principles of a phased array, mismatches in pulse intensity will result in non-uniform destructive interference along non-steered directions. Second, highlighted by the asymmetric nature of the 4-direction beam steering shown in Fig. 7, the construction of the composite thermoelastic acoustic transducer led to inhomogeneities in optical absorption. While the circular illumination pattern (phase ramp) irradiated the same location on the transducer surface, the rotations in the azimuthal direction that is necessary for beam steering in different directions led to the four beam patterns falling on different regions of the circular area on the transducer surface. Furthermore, as shown in Fig. 9, the profiles of the acoustic pulse depend upon the number of combined hologram components that construct it –namely by increasing the period of the resulting pulse. This differentially affects the propagation of each composite pulse thus leading to nonuniform interference. Lastly, the surface of the transducer is not perfectly flat, and therefore the height variations manifest as phase variations leading to further deviations from ideal interference.

While a time delay can be accomplished via fiber coupling, this investigation chose to showcase the abilities of free-space optical delay. First, as is common when dealing with high power densities, the risk of laser induced damage is great concern –one which is mitigated in a free-space design. An obvious trade-off related to overall optical train alignment stability is apparent, however there exists a nonnegligible freedom in pointing tolerance provided the vibrational and thermal environment is calm and invariable. A free-space design further permits easy access to beam modification at numerous points along the optical path. While not demonstrated in this study, intra-path beam modification can prove to be beneficial to maintain homogeneity in intensity, diameter, polarization, etc. Finally, a free-space design allows for straightforward tuning of the optical delay line without the need for new hardware. Realignment of the aforementioned optical delay apparatus is not trivial, however it is considerably less involved than replacing optics, fibers, hardware, etc.

**Table 1.** Measured beam steering with comparison to simulation. For each of the cardinal directions shown in this study, the (x, y) deviations in mm are displayed, with boldface numbers signifying intended direction.

| Method | Total Beam Shift (x, y) [mm] |  |  |  |
| --- | --- | --- | --- | --- |
|  | Down | Up | Left | Right |
| Simulation | (0.0, **-2.1**) | (0.0, **2.1**) | (**-2.1**, 0.0) | (**2.1**, 0.0) |
| Measurement | (0.4, **-2.1**) | (0.4, **1.9**) | (**-0.9**, 0.1) | (**1.9**, 0.2) |

### 5.2. Towards an Arbitrary Acoustic Wavefront

The overall extent to which beam steering can be accomplished by the phased array technique laid out in this study is largely dependent upon the amount of total photoacoustic phase delay available. While this study served as a demonstration of a wavefront generation technique by constructing a simple phase ramp, the possible extensions of the technique are far reaching. Indeed, with enough phase-delayed components, arbitrary phase maps can be constructed resulting in wavefront control analogous to optical techniques [55, 56].

Prior testing of phase wrapping across the acoustic aperture, namely when utilizing a single oscillation, has shown to produce considerable artifacts and side lobes. Increasing the available phase delay to incorporate multiple oscillation cycles can foster even more flexibility in wavefront generation. These advancements are practical concerns, however, and are not theoretical limitations. Indeed, extensions of the method described in this study can yield conceptually arbitrary wavefront generation.

### 5.3. Concluding Remarks

A photoacoustic beam steering technique has been demonstrated which uses a free-space optical beam delay and SLM-generated holograms to produce an optically-driven acoustic phase ramp on a thermoelastic polymer. The resulting acoustic field was measured at a propagation distance of 20 mm with a steering angle of around 5 degrees, in agreement with Rayleigh-Sommerfeld diffraction principles. The acoustic aperture is highly configurable as is the related phase delay facilitated by the optical delay line. This serves as a verification of a new photoacoustic wave-front generation process, which allows for theoretically arbitrary acoustic phase profiles.

## Acknowledgments

CJR acknowledges funding from EPSRC (EP/S016538/1, EP/X017842/1 and EP/W024969/1), BBSRC (BB/T011947/1 and BB/X004716/1), Wellcome Trust (212490/Z/18/Z), Cancer Research UK (29694 and EDDPMA-May22*\*100059), the Royal Society (RGS*\*R2*\*212305 and IES*\*R2*\*222231), the Chan Zuckerberg Initiative (2020-225443 and 2020-225707) and the Imperial College Excellence Fund for Frontier Research.

## Supplementary Material

A render of the beam delay design is shown in Fig. 13a. The small mirrors in the octagonal arrangement are relayed to other small mirrors in succession by means of the larger mirrors. Top-down beam paths for the first two passes are shown in Figs. 13b and 13c. The green highlight shows the start of each pass while the red highlight shows the end, with some light escaping during each pass at the small, partially-reflective mirrors in the center. While the device can utilize a total of eight partially-reflecting mirrors, this study used only four beam components. The partially reflecting mirrors consist of a single Thorlabs BST05 (70:30 R:T) and two Thorlabs BSW05 (50:50 R:T). Considering the reflection percentages of the partially-reflective mirrors along with optical loses on each surface, the resulting pulse intensities at the output of the device are roughly 30%, 21%, 24%, and 24% of the initial pulse, respectively. Seeing as the use of an SLM can regulate the intensities of the four pulses independently in numerous ways (diffraction efficiency, coupling, etc.), this was taken as an acceptable first-order intensity regulation method.

**Figure 13:**
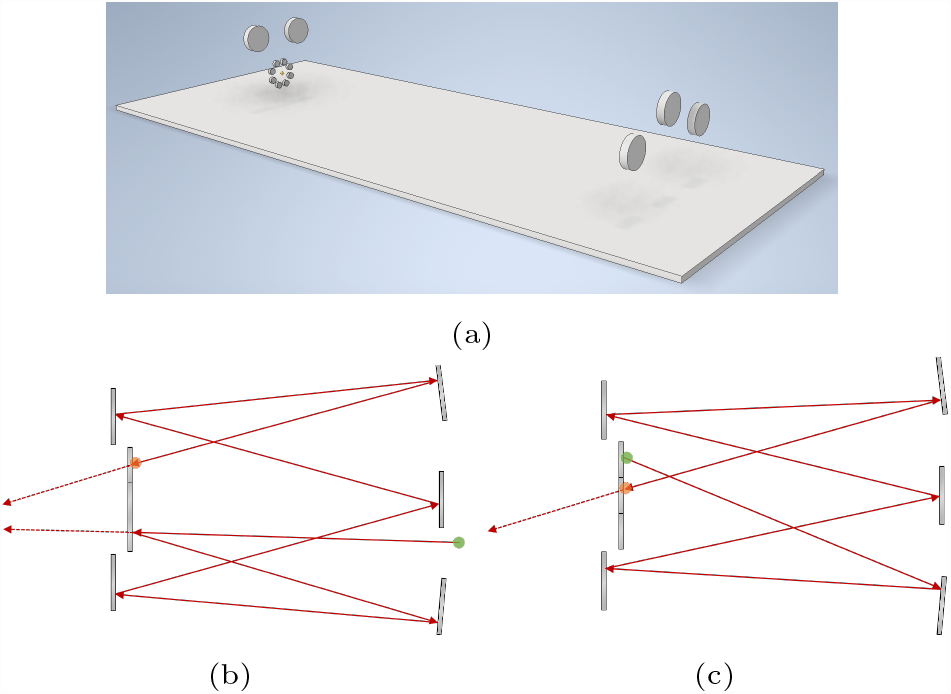
The custom optical delay design. **(a)** shows a rendering of the optics (mounts and supports not shown), while **(b)** and **(c)** show the first two beam path loops, respectively. The start of each loop is highlighted in green while the end is highlighted in red. The small mirrors on the left side are partially-reflective, allowing some light to escape on each pass. This design is more configurable than other multipass cells in that the mirrors can be translated and realigned to modify the total time delay.

